# A general framework to learn tertiary structure for protein sequence characterization

**DOI:** 10.1101/2021.04.01.438098

**Authors:** Mu Gao, Jeffrey Skolnick

**Affiliations:** Center for the Study of Systems Biology, School of Biological Sciences, Georgia Institute of Technology, Atlanta GA 30332, USA

**Keywords:** protein structure prediction, protein folding, sequence alignment, deep-learning, structural alignment

## Abstract

During the past five years, deep-learning algorithms have enabled ground-breaking progress towards the prediction of tertiary structure from a protein sequence. Very recently, we developed SAdLSA, a new computational algorithm for protein sequence comparison via deep-learning of protein structural alignments. SAdLSA shows significant improvement over established sequence alignment methods. In this contribution, we show that SAdLSA provides a general machine-learning framework for structurally characterizing protein sequences. By aligning a protein sequence against itself, SAdLSA generates a fold distogram for the input sequence, including challenging cases whose structural folds were not present in the training set. About 70% of the predicted distograms are statistically significant. Although at present the accuracy of the intra-sequence distogram predicted by SAdLSA self-alignment is not as good as deep-learning algorithms specifically trained for distogram prediction, it is remarkable that the prediction of single protein structures is encoded by an algorithm that learns ensembles of pairwise structural comparisons, without being explicitly trained to recognize individual structural folds. As such, SAdLSA can not only predict protein folds for individual sequences, but also detects subtle, yet significant, structural relationships between multiple protein sequences using the same deep-learning neural network. The former reduces to a special case in this general framework for protein sequence annotation.

## 1 Introduction

The amino acid sequence of a protein encodes the information for carrying out its function. One essential aspect is the tertiary structure of the protein. Indeed, the prediction of protein tertiary structure from its sequence is a fundamental question in biophysics (1). In order to predict protein structure at high accuracy, one main challenge is to model the long-range, many-body effects that collectively dictate a protein’s tertiary structure (2). Over the past several years, exciting breakthroughs have been made to better address these long-range interactions (2). Using a deep residual convolutional neural network, significant success has been demonstrated in predicting contacts between individual residues of a protein sequence (3). Such residue-residue contacts yield both the local secondary structure and the global fold, and it is the accurate prediction of their synergy that improves model quality. Subsequently, several groups demonstrated that better residue-residue contact or detailed distance matrix (distogram) predictions led to significantly improved structure predictions, especially for challenging targets (4–8). In CASP13, a blind biannual protein structure prediction competition, all four top-ranked groups in the most challenging, free-modeling category used residue-residue contacts or distance matrices predicted via deep-learning (9). Among them, DeepMind’s AlphaFold achieved the best performance using a high-quality distogram to derive statistical folding potentials (6). In CASP14, many improved deep-learning approaches using the convolutional residual networks were presented, e.g., (10), but AlphaFold2 dominated the competition using a new, end-to-end deep-learning algorithm with an attention mechanism (11).

A topic closely related to protein structure prediction is protein sequence comparison or alignment (12). In the low pairwise sequence identity regime of less than 30%, two protein sequences may exhibit no apparent sequence similarity yet display significant fold similarity when their structures are revealed and superimposed (13). This observation is attributed to the fact that the structural space of protein folds is very small for both evolutionary (14) and physical reasons (15). Traditionally, a variety of sequence alignment approaches have been developed and applied to assist protein structure prediction, e.g., Hidden Markov Model (HMM) (16) and “threading” approaches (17–19). These efforts provide the foundation for template-based modeling approaches (20). Conversely, if the structures encoded in the two sequences are known, their structural alignment generally leads to a more accurate sequence alignment than those from classical sequence alignment approaches. Such an accurate, meaningful alignment is often the key to understanding what a novel protein sequence does, e.g., predicting functional sites (21, 22).

Naturally, this leads to a question: can deep-learning be directly applied to generate a protein sequence alignment with an accuracy close to the structural alignment counterpart? If so, this would not only extend the ability to recognize evolutionarily distant sequence relationships but also enable a deeper learning of the folding code. Moreover, it has practical applications for function and structure prediction and possibly evolutionary inference. To answer this question, we recently developed SAdLSA, a sequence alignment algorithm that uses a deep convolutional neural network to learn many thousands of structural alignments (23). The advantage of SAdLSA was demonstrated in benchmark tests against HMM-based HHsearch (24). For challenging cases, SAdLSA is ~150% more accurate at generating pairwise alignments and ~50% more accurate at identifying the proteins with the best alignments in a sequence library. This allowed the program to detect remote relationships that may be useful for genome annotation or functional predictions.

Given the encouraging benchmarking results of SAdLSA, one would like to understand why it performs better than classical sequence comparison approaches. Obviously, the deep-learning algorithm plays a key role in this improvement, but how does it work? Previously, we have speculated that SAdLSA implicitly learns the protein folding code without offering direct evidence. In this study, we shall further substantiate this claim. We noticed that when the same sequence was input into SAdLSA, SAdLSA aligns the sequence against itself, i.e., self-alignment, and outputs an intra-sequence distogram for the input. This distogram could encode the fold much like a deep-learning algorithm designed to predict the distogram for a single query sequence, e.g., DESTINI (4). We shall perform analysis to understand the distogram generated by SAdLSA self-alignment and demonstrate that SAdLSA provides a more general framework to learn protein structures for sequence annotation purposes.

## 2 Methods

For this study, we mainly employ SAdLSA, a deep-learning (DL) based approach for protein sequence alignment. The details of SAdLSA and the benchmark results have been described in detail elsewhere (23). Here, we briefly recapitulate its key features.

### 2.1 SAdLSA

An overview of SAdLSA is presented in Figure 1. The inputs to this network are two position-specific sequence profiles, each of dimension *N_k_* × 20, where *N_k_* is the length of the *k*-th sequence (*k* = 1, 2), and the 20 columns represent the 20 different amino acids at each residue position (hence position-specific). Here, we use the profiles generated from HHblits (25). In machine-learning language, the sequence profiles are also known as embeddings. The outer product of these two 1D sequence features yields a 2D matrix of features, where at position (*i,j*) of the matrix the elements are a concatenation of the 20 columns formed from the *i*-th residue of sequence 1 and the *j*-th residue of sequence 2. Subsequently, these 2D features are fed into a fully convolutional neural network consisting of up to 34 residual blocks. The main objective of this neural network model is to predict residue-residue distances between the two input sequences that recapitulates their optimal structural alignment, using observed structural alignments as the training ground truth. The training distance labels are created from structural alignments by APoc (21), which takes advantage of a global alignment provide by TM-align (26).

**Figure 1.**
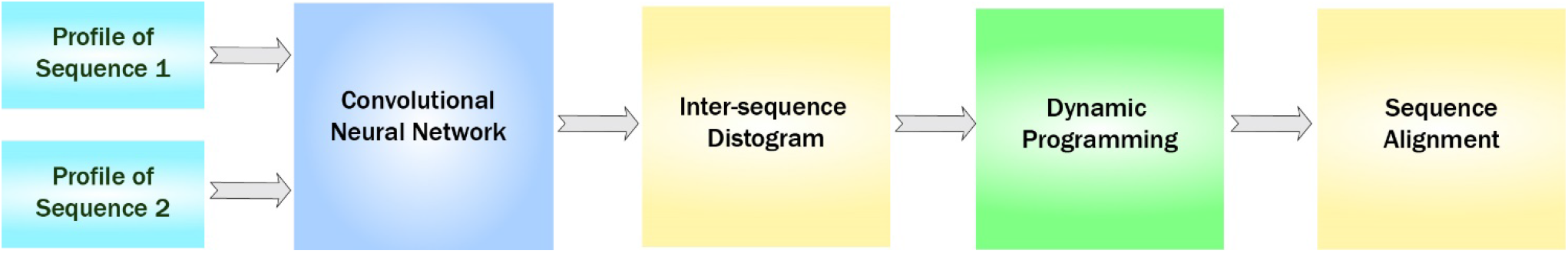
Flowchart of SAdLSA, a deep-learning algorithm for protein sequence alignment. In this study, the same sequence input is supplied to the SAdLSA pipeline, resulting in an intra-sequence C_α_-C_α_ distogram prediction of a single protein, instead of a typical application scenario, whereby the inter-sequence distogram portraying the superimposition of two different proteins is predicted and utilized for deriving their sequence alignment using dynamic programming.

The neural network is composed of multiple residual blocks, either conventional (27) or dilated (28) in slightly different design variants. The residual block design is a key to train a deep neural network model. Within a residual block, each convolutional layer is composed of about 64 filters with a kernel size of 3×3 or 5×5. After the residual blocks, the last 2D convolutional layer outputs 22 channels, representing 21 distance bins (1 to 20 at 1 Å each, and >20 Å) and channel 0 which is reserved for ignored pairs (e.g., gap residues missing in a structure, or large distances >30 Å). Finally, a softmax layer calculates the probability scores for each distance bin. Here, the same network was also applied to the same two input. The mean probability scores of these two runs are the final output score for this DL model. This ensures that the alignment is identical if one swapped the position of two input sequences. For this study, we used the consensus scores from six DL models, including three models with 14 residual blocks and 64 5×5 kernels in each convolutional layer, and three dilated model with 34 residual blocks (alternating 1,2,8,16,32 dilate rates) and 50 to 75 3×3 kernels. The two type of DL models have 2.9 and 2.4 million parameters, respectively.

The outputs from a DL model are the probabilities of distance bins forming an inter-protein residue-residue distance matrix. To build an alignment using dynamic programming (DP), we convert this probability matrix into a mean distance matrix *D*, whose element

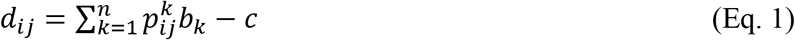

where *i,j* are target/template sequence positions, 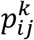 is the probability for bin *k* at position (*i, j*), *b_k_* are distance labels from the sequence (1, 2, … 20, 22). *D* is subsequently adapted as the scoring matrix to obtain the optimal alignment using a Smith-Waterman-like DP algorithm (29). The distance matrix *D* is also used to calculate an estimated TM-score (30, 31) for ranking the significance of an alignment. The constant *c* is set at 1 such that a perfect alignment gives an estimated TM-score of 1.

Since we study the self-alignment of a given sequence, we feed exactly the same sequence profiles into SAdLSA, and the resulting sequence alignment itself is universally at 100% identity, with a predicted TM-score ~1. We focus on the residue-residue distance matrix *D*, which is converted from a general inter-sequence scenario into a special intra-sequence scenario, because the two input sequences are the same.

### 2.2 Data sets

We employed the same test and training sets from the original SAdLSA study (23). Both sets are curated from the SCOP database (32). The training set is composed of 79,000 pairs from a SCOP30 set of ~5000 domains. These training protein domain sequences share less than 30% identity. The test set is an extrinsic test set of the sequences of 593 randomly selected protein domains from 391 SCOP folds. Homologs of the testing sequences at 30% sequence identity or higher, or with a BLAST E-value < 0.1, are excluded from the training set. In this study, we employed SAdLSA models trained on this training set and conducted SAdLSA self-alignments on each of the 593 test sequences.

### 2.3 Analysis

#### 2.3.1 Distogram assessment

It has been well-established that residue-residue contacts characterize a protein structural fold (2, 4). The most common definition of a protein contact is based on the distance between C_α_ or C_β_ atoms. That is, a contact between a pair of protein residues *i* and *j* is defined if the Euclidean distance between their C_α_ (or C_β_) atoms is less than a cutoff value, typically at 8 Å. A popular contact metric adopted by the CASP assessors (33) is the precision of the top *L*/*k* C_β_–C_β_ contact predictions within short, medium or long ranges, where *L* is the length of the target and *k* = 1, 2, and 5, and the sequential distance *s_ij_* of residues *i* and *j, s_ij_* ≡ |*i* – *j*|, defines the range: *short* (*s_ij_* ∈ [6, 11]), *medium* (*s_ij_* ∈ [12, 23]), and *long* (*s_ij_* ∈ [24, ∞)), i.e., nonlocal residue pairs. Since SAdLSA was trained on the distances between C_α_ atoms, we use C_α_–C_α_ contacts with an 8 Å cutoff as our definition of protein inter-residue contacts and consider only those belonging in either the medium- or long-range regime, i.e., *s_ij_* ∈ [12, ∞). The predictions are ranked by the probability of forming a C_α_–C_α_ contact. To obtain the probability, one simply sums the probabilities for distance bins from 0 to 8 Å, since the SAdLSA DL models output a probability matrix *D* for 21 distance bins. This probability score is then employed for the precision analysis as outlined above. The precision is defined as TP/(TP+FP), where TP is the number of true positives, i.e., native contact observed in an experimental structure, and FP is the number of false positives within the top *L/k* contact predictions evaluated.

In addition, we introduce the Mean Absolute Error (MAE) of the predicted distogram versus the ground truth distogram. Specifically, we calculate the MAE using the coordinates of the C_α_ atoms of nonlocal residue pairs, i.e., the sequential distance of residue pairs is no less than 6. The overall MAE for a distogram is defined as

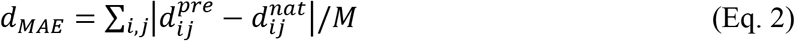

where (*ij*) are the indexes of a pair of nonlocal residues separated up to 20 Å in the native distogram, *M* the total of such pairs, 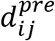 the predicted distance by the SAdLSA self-alignment, and 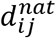 the distance observed in the native distogram. Additionally, for each target, we also obtain the MAE values within each distance bin from 4 to 20 Å. If a target does not have any nonlocal residue pair within a specific 1Å bin, the MAE calculation is skipped for this bin.

In order to estimate the statistical significance of *d_MAE_*, we consider its expected value using a background distribution based on the pairwise residue distances observed in the training distograms. For each distance bin, we count the residue pairs within this bin as observed in the training ground truth distograms, and then obtain the observed frequency *f^k^* for this distance bin by dividing the count against all counts of all bins from 1 to 21 (inter-residue distances between 20 to 30 Å are used for bin 21 as they are what collected for training). Substituting *f^k^* for 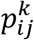 in Eq. 1, we obtain the expected value of *d_ij_*, 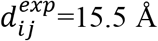 Å for any (*i,j*) according to this reference distribution. Note that our training ground truth distograms are of inter-sequence residues, in comparison to the distograms of intra-sequence residues from the self-alignment. But, since the inter-sequence distograms are actually employed for SAdLSA training, it is appropriate to use the distance distribution collected from these distograms as the reference background. This leads to a naïve way to calculate the expect *d_MAE_*, 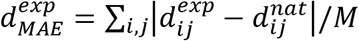, yielding a mean 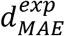 of 3.07 Å and a standard deviation of 0.20 Å by applying the formula to 4,661 structures employed in the training set. Likewise, for comparison the same formula is applied to the test set, including the overall error and errors for individual distance bins. Using the one-tailed test for a normal distribution, one can calculate the *p*-value of an *d_MAE_* value using the background distribution derived from the training set.

#### 2.3.2 t-SNE analysis

We performed t-distributed Stochastic Neighbor Embedding (t-SNE) (34) to analyze the factors that contribute to the precision of contact predictions by SAdLSA self-alignment. This is a nonlinear dimensionality reduction tool more suitable for this analysis than a classical principal component analysis. Five features were used for this analysis including (i) the best training pair for each target measured by their TM-score (33), (ii) the ratio of the total observed native contacts (combining both medium- and long-range contacts, i.e., *s_ij_* ∈ [12, ∞)) over the sequence length, whether the best training pair belongs to (iii) the same fold or just the (iv) superfamily as the target according to the SCOP classification, and (v) the sequence diversity of the multiple sequence alignment of the target. The TM-score is a protein length-independent metric ranging from 0 to 1, and a TM-score > 0.4 indicates a statistically significant alignment (26). We use (0,1) to represent the logical variables (e.g., is it a member of the same SCOP fold or not). The sequence diversity is calculated with the Multiple Sequence Alignment (MSA) of the target and defined as 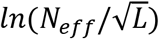 taken from (23), where *N_eff_* is the number of effective sequences in the MSA without length normalization. In our t-SNE analysis, we employed the default parameters including the perplexity parameter set at 30.

### 2.4 DESTINI2

For comparison purposes, we employed DESTINI2 to conduct inter-residue distance prediction on the same test data set, i.e., 593 SCOP domain sequences, used to benchmark SAdLSA. DESTINI2 improves DESTINI by using a deeper, dilated convolutional residual network model. In this study, we used 39 dilated residual blocks similar to the one implemented in SAdLSA. For training, about 10000 crystal structures with < 2.5 Å resolution were taken from a March 2020 release of PISCES (35), which were curated from the PDB database (36). For a fair comparison, we retrained DESTINI2 models by removing the test set entries and their close homologs from the original DESTINI2 training set using the same criteria as that used for the SAdLSA test.

## 3 Results

We conducted self-alignments for 593 target sequences with SAdLSA, using deep-learning models trained without obvious homologs to any of the 593 target sequences (see Methods). As one would expect, SAdLSA returns a sequence alignment at 100% identity and a predicted TM-score close to 1. This seems trivial. But, if one carefully inspects the residue-residue distogram prediction, it not only contains information giving rise the identical alignment, but also contains inter-residue distance information to structurally characterize the fold encoded by the sequence itself.

### 3.1 SAdLSA self-alignment generates a fold-depicting distogram

Figure 2 illustrates an example of a SAdLSA self-alignment prediction. This 205 AA target sequence encodes a classic Rossman fold, which is composed of an βαβ alternating secondary structural segments found in many nucleotide-binding proteins (37). The characteristics of this fold are displayed in the distance plot calculated between the C_α_ atoms determined in the crystal structure (Figure 2, top left panel). The remaining residue-residue distance plots are generated by the SAdLSA self-alignment of the same sequence. These are from the 21 scoring channels designed to predict the probability of each pair of C_α_ atoms falling into each distance bin from 0 to beyond 20 Å. The first three channels are straight diagonal lines, which give rise the 100% identity in the sequence alignment and are not the focus of this study. Starting from plots ≥ 4 Å, one recognizes inter-residue distance relationships. First, the immediate neighboring residues are shown between 3-4 Å. Then, the main secondary structure elements including the fold’s six α-helices and seven β-strands exhibit their patterns in the 4-5 and 5-6 Å plots. The packing between these secondary structural elements becomes clear in the subsequent three plots from 6-9 Å. The detailed packing orientations among secondary structural elements are further delineated in the remaining distance plots up to 20 Å. Finally, the highlights in the >20 Å plot signal the regions that are distant from each other. The top *L* medium- or long-range contact predictions for this case have a precision of 86%, which is sufficient to reconstruct a high-resolution structural model whose TM-score > 0.7 (2). The mean absolute error *d_MAE_*, calculated from the C_α_–C_α_ distances of all nonlocal residue pairs separated up to 20 Å, is only 1.57 Å from the native distogram.

**Figure 2.**
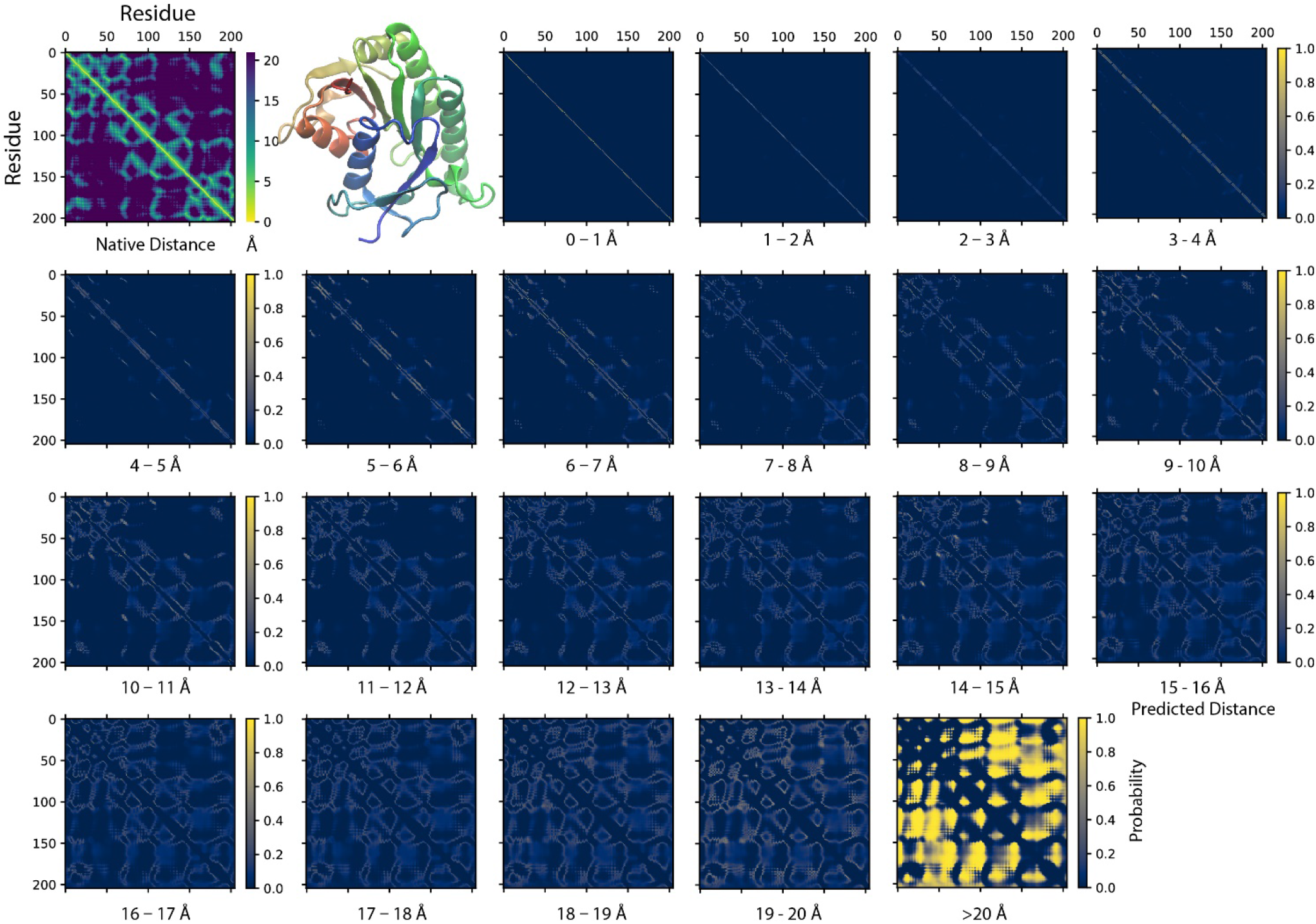
An example of predicting residue-residue distance matrices by SAdLSA self-alignment. The native (experimental) structure (PBD ID: 1hdo, chain A; SCOP ID: d1hdoa_) of the target sequence, human Biliverdin IXβ reductase, encodes a classic Rossman fold shown in the cartoon representation using the red-green-blue color scheme from the N- to C-terminus. For each pair of residues, its experimentally observed (native) C_α_-C_α_ distance is shown in the upper left corner. The remaining 21 probability density plots are generated by SAdLSA self-alignment for the target sequence. Each plot predicts the probability of C_α_-C_α_ distance within a distance bin from 0 to 20 Å at 1 Å spacing, and the probability > 20 Å in the last plot.

How accurate is the SAdLSA self-alignment for predicting residue-residue distances in general? If one examines the *d_MAE_*, the overall number looks good with a mean of 2.43 Å, and 92.7% of targets are below 3 Å (Figure 3 insert). By comparison, if one naively assigns distance distribution according to the observed fractions from the training set (see Methods), one obtains a mean expected error, 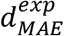, at 3.08 Å. Overall, 96.6% of targets have a smaller distance prediction error by SAdLSA than the value from the naïve reference approach. Moreover, about 408 (69%) targets have a significant *d_MAE_* below the *p*-value cutoff of 0.05. Figure 3 further details the distributions of MAE between the SAdLSA predicted and the corresponding native distograms for individual distance bins from 4 to 20 Å. It is clear that residues forming direct contacts within the first five bins are most challenging to predict and exhibit large variations, with the mean MAE gradually decreasing from 6.6 Å in the 3-4 Å bin to 3.4 Å in the 8-9 Å bin. But these distance predictions by SAdLSA are actually highly significant in comparison to the expected values whose MAE errors are up to 8 Å higher on average than SAdLSA predictions. On the other hand, the large distance bins from 14 to 18 Å yield relatively small MAE values < 2.5 Å, but it is not surprising as the expected MAE is below 2 Å.

**Figure 3.**
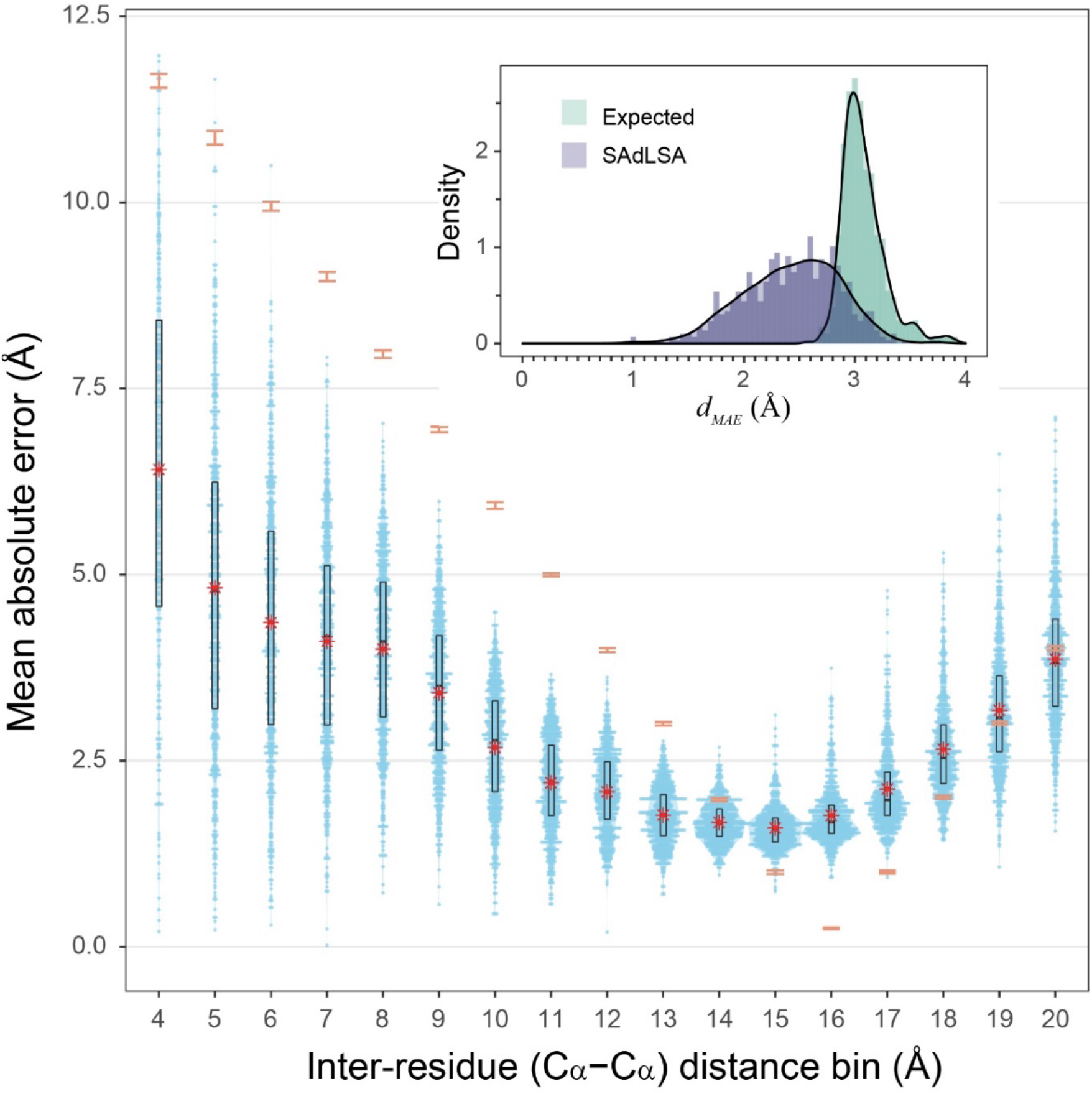
Mean absolute error of predicted distogram *vs* native distogram for 593 test cases. The main plot displays the distribution of MAE values within each C_α_-C_α_ distance bin from 4 to 20 Å for nonlocal residue pairs. Each bin has a spacing of 1 Å and each blue point represents a target protein. Violin contours are proportional to the counts of targets at different MAE levels with a bin width of 0.025. Black boxes and bars represent the 2^nd^ and 3^rd^ quartiles (25% to 75% ranked by MAE values) and the median of the distributions. The red stars represent the mean values. For comparison, the expected MAE distributions are shown in orange error bars, which are centered at the mean and extended to ± sd. The insert shows the histograms of the *d_MAE_* values in two separate assessments, calculated (purple) from the SAdLSA self-alignment and expected (teal) from the background distribution. For each target, its *d_MAE_* value is calculated from all nonlocal residue pairs up to 20 Å as observed in the native distogram.

From a structural perspective, direct inter-residue contacts are the most important. Moreover, one only needs to predict a fraction of the total number of contacts in order to obtain a correct fold prediction (4). We therefore turn our focus to inter-residue contacts. According to the benchmark results on 593 protein sequences, the mean precisions are 40.3%, 52.6% and 63.7% for the top *L, L/2, L/5* predictions of medium- or long-range inter-residue C_α_ contacts, respectively (Table 1). The detailed distributions of individual predictions are shown in violin plots (Figure 4). For instance, in the middle top *L*/2 plot, about half of the sequences have a precision value > 50%, which is sufficient to derive the correct fold for a single-domain using these predictions (4). At *L*/5, 107 (18%) entries have 100% precision, and the median precision is at 69%. These numbers are not as good as the deep-learning approaches specifically trained to predict residue distograms, e.g., they are about 25% worse than DESTINI2 (Table 1). Given the fact that SAdLSA was not trained to predict residue distances for a single structure (see more reasoning in Discussions), but rather to recognize the similarity between pairs of structures, these results are encouraging and clearly demonstrate the generality of the SAdLSA framework for learning protein structures.

**Figure 4.**
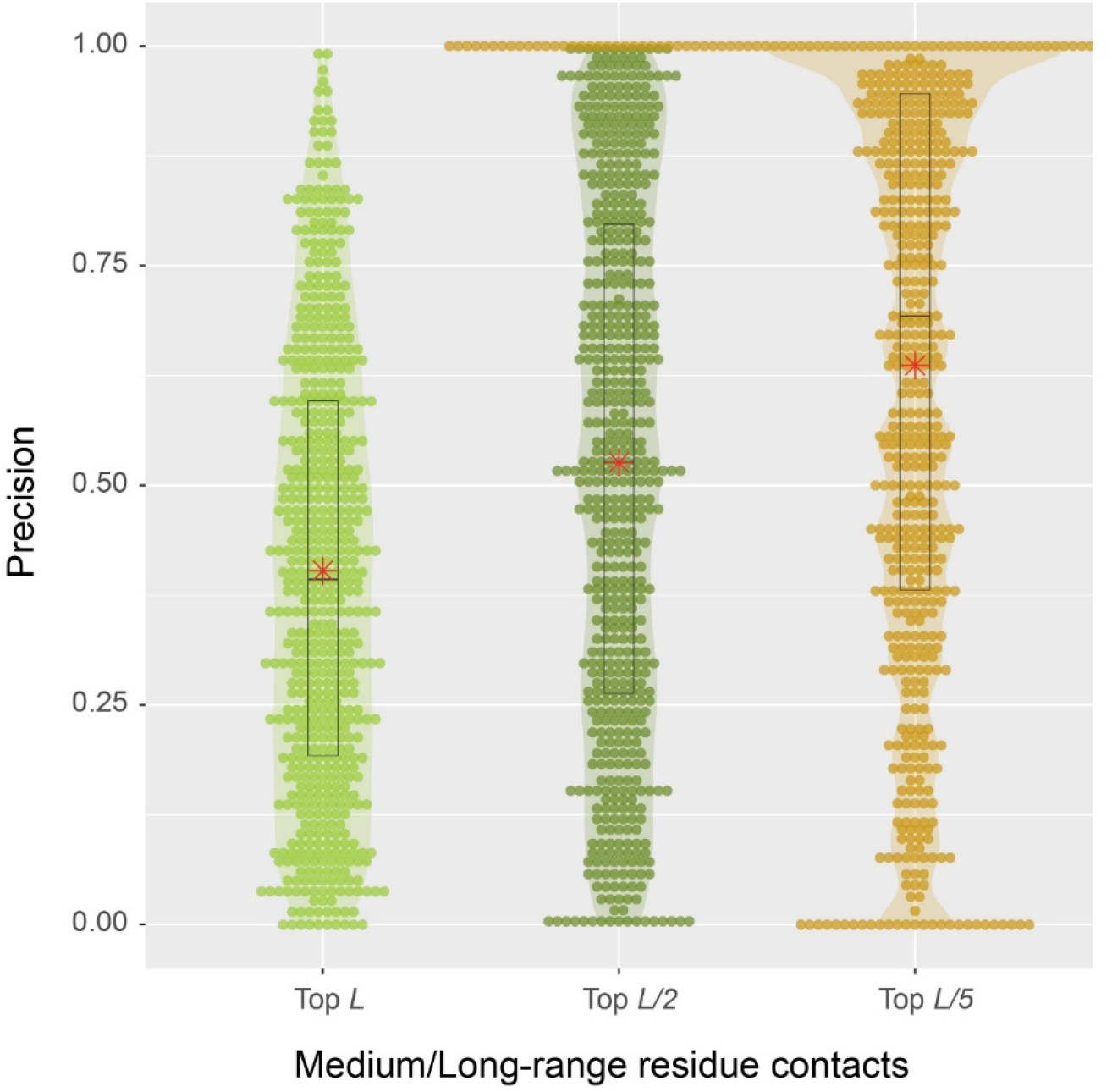
Precision of interresidue contact predictions via SAdLSA self-alignment. The precision of the top *L* (light green), *L*/2 (green), and *L*/5 (gold) are shown as circles for each target sequence. Violin contours are proportional to the counts of targets at different precision levels with a bin width of 0.01. Black boxes, median bars and means are represented following the same plotting scheme as in Figure 3.

**Table 1.** Mean precision of Medium/Long-range inter-residue contacts for 593 targets.

| Method | Medium/Long-range Contacts* |  |  |
| --- | --- | --- | --- |
| | $L$ | $L/2$ | $L/5$ |
| SAdLSA ($C_{\alpha} - C_{\alpha}$) | 0.403 | 0.526 | 0.637 |
| DESTINI2 ($C_{\alpha} - C_{\alpha}$) | 0.645 | 0.777 | 0.858 |
| DESTINI2 ($C_{\beta} - C_{\beta}$) | 0.678 | 0.803 | 0.879 |

\* Medium/Long-range Contacts denote residue-residue contacts whose sequential distance are in either the medium or long-range regime. Contact predictions are converted from the distogram predicted by each method. All deep-learning models were re-trained to exclude the test set and their close homologs from the original training sets (see Methods).

### 3.2 What contributes to successful distogram prediction by SAdLSA self-alignment?

Next, we seek to understand the factors that affect the accuracy of distogram predictions by SAdLSA self-alignment. Like all machine-learning algorithms, the capability of distogram predictions must come from the training data of SAdLSA. Although SAdLSA was not trained to learn individual protein folds, we hypothesize that the fold-depicting distogram provided in the SAdLSA self-alignment comes from learning the training pairs sharing fold similarity as the target sequence. During SAdLSA training, it mainly focuses on aligned residues across two different sequences, but the network also observes the relative positions among aligned residues. As a result, SAdLSA learns the folding code of individual protein sequences, provided that they exhibit significant fold similarity. The more similar in their structures, ideally the same fold as the training pair, the more likely the distogram pattern is learned for this specific fold.

#### 3.2.1 Presence of the same fold as a target in the training set is important

To explore the validity of this hypothesis, for a target structure and a pair of training structures, we introduce *T*, defined as the minimal TM-score of the two training structures with respect to the target structure, using the target to normalize the TM-score. For each target, we find the highest *T*, denoted as *T*\*, among all pairs in the training set. Figure 5 shows the correlations between *T*\* and the precision for the top *L*/2 residue-residue contact predictions. The violin plots demonstrate a clear upward trend in precision from the low *T*\* regime to high *T*\* regime. When there is a lack of obvious training structures that are similar to the target, e.g., when *T*\* < 0.4, the median precision is only at 39%. The same metric increases to 52% if *T*\* ∈ [0.5, 0.6), and dramatically to 82% if *T*\* > 0.7. The Pearson correlation coefficient between *T*\* and the precision of all targets is 0.34, which clearly shows the dependency of precision on *T*\*, despite the indication that other factors are also in play.

**Figure 5.**
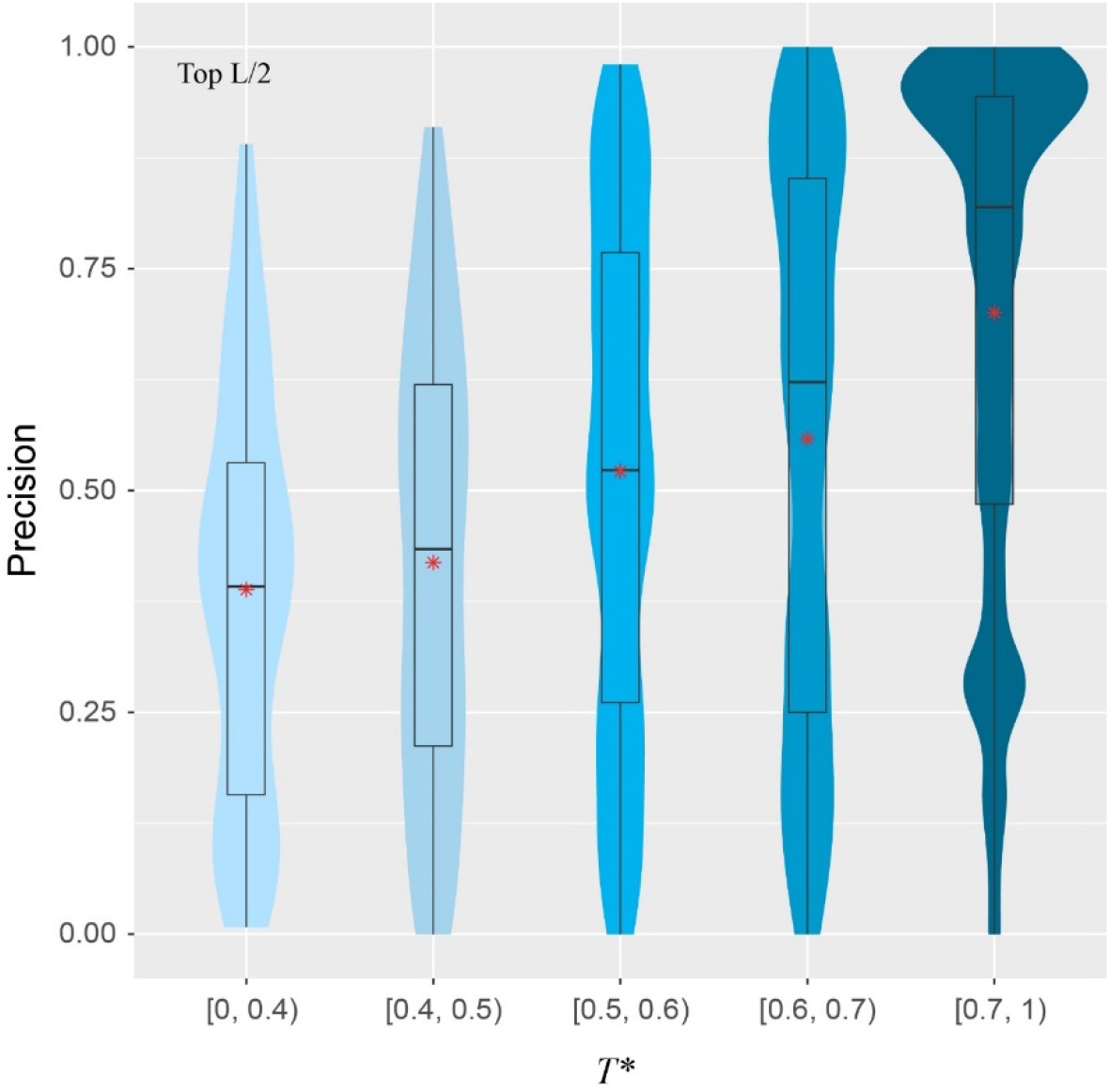
Precision of inter-residue contact prediction *vs T*\*, the highest TM-score to all structures in the pairs found in the training set. The violin and box plots follow the same scheme as in Figure 3.

For example, Figure 5 shows that there are still very good distogram predictions in the low *T*\* regime. How could SAdLSA accurately predict a distogram when there is no similar training structure? There are 11 target sequences with *L*/2 precision > 60% within *T*\* < 0.4. If one examines these structures, some of them are composed of multiple domains or subdomain structures. For example, one target, with SCOP ID d3u7qb, is a 522 AA sequence composed of three Rossman folds and two helical bundle domains, despite the fact that SCOP defines it as a single domain. Although there is no other structure in the training set that resembles the overall structure of this target, the folds of its individual domains can be learned separately. As shown above, the Rossman fold is relatively easy to learn (Figure 2). As a result, the overall precision prediction is 76% and the *d_MAE_* is 2.44 Å (*p*-value = 8.0×10^−4^) for this query sequence. More interesting examples are analyzed below.

#### 3.2.2 Evolutionary relationships facilitate fold-learning

What are the other possible contributing factors? In addition to *T*\*, we further consider four other features including the ratio of total observed native contacts (within either the medium- or long-range regime) over the sequence length, whether the *T*\* training pair belong to the same fold or superfamily according to the SCOP classification, and the sequence diversity of the multiple sequence alignment of the target. The Pearson correlation coefficient between each of these four additional features and the contact precision is 0.55, 0.30, 0.33, and 0.33, respectively. Figure 6 shows the results from t-SNE analysis of these five features. Three big clusters emerge and represent targets whose *T*\* training pair are from the same SCOP superfamily (240 entries), the same SCOP fold but not the same superfamily (40 entries), and different SCOP fold (313 entries). They exhibit different levels of difficulty for contact prediction, at mean precision values of 64%, 60%, and 43%, respectively. This result makes sense as the sequence profiles from the same superfamily are similar and relatively easy to learn for a neural network model, whereas sequence profiles from remote families or those without an apparent evolutionary relationship are much harder to learn. In the same superfamily cluster, there are very few “bad” predictions, e.g., 41 (17%) targets at *L*/2 have a precision < 30%. One explanation is the structural variation between the target and the training structures, despite the fact that they are from the same superfamily. On the other hand, even for cases where the evolutionary relationship might not be clear, it is still possible to predict a fold-depicting distogram reasonably well with SAdLSA, as is evident in the clusters where the SCOP superfamilies are different. In particular, we note that SAdLSA performs very well for a target structural fold if there are training structures originated from the same SCOP fold but not necessarily the same SCOP superfamilies, as is evident in the mean precision of 60% among 40 such entries, close to the value of 64% in the cluster representing the same superfamily.

**Figure 6.**
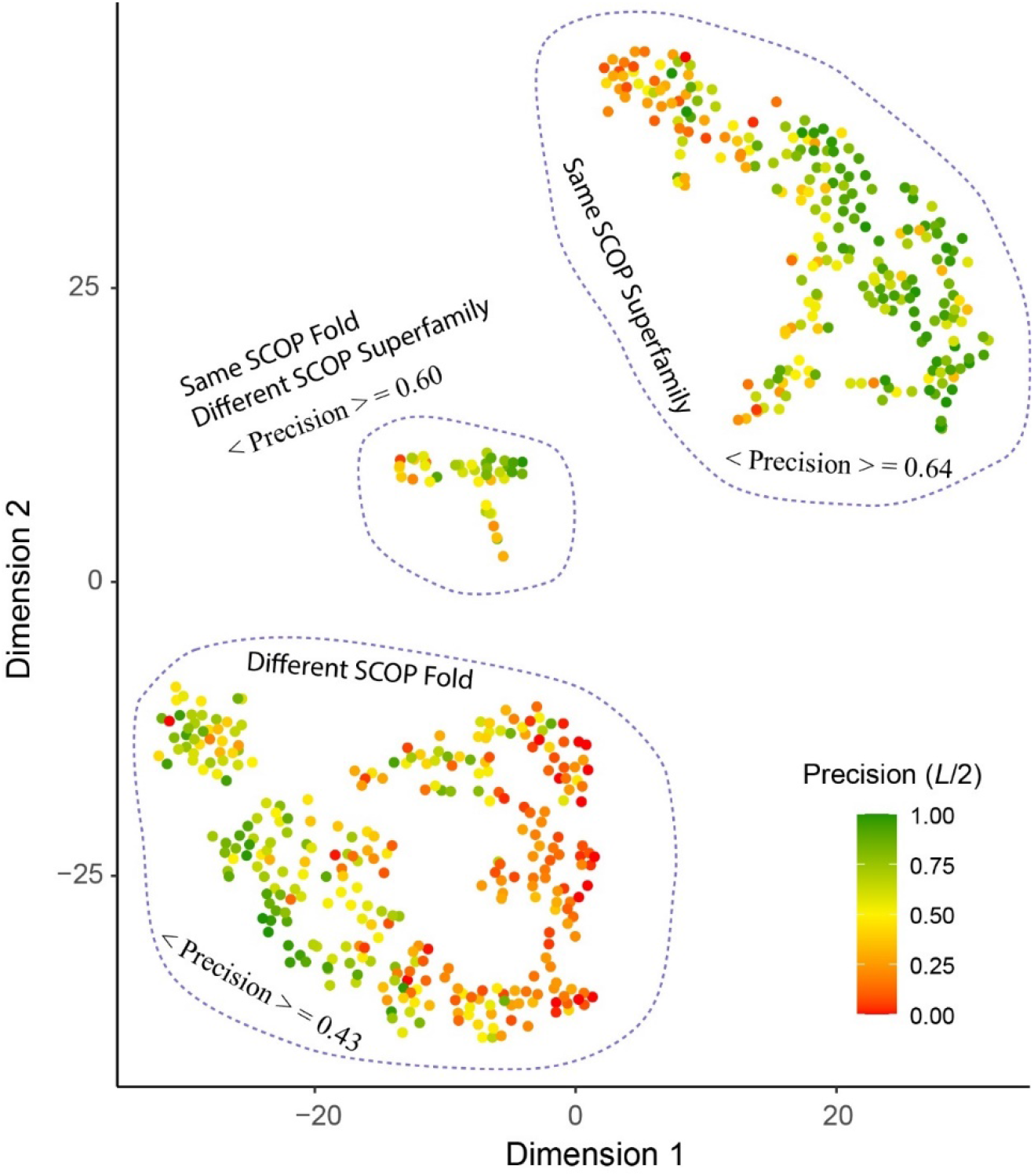
T-SNE analysis of factors affect the precision of inter-residue distance predictions. Each point in the plot represents one of 593 target sequence, color-coded according to its precision value of the top *L*/2 Cα-Cα contact prediction for those belonging in the medium- or long-range regime. The template pairs structurally most close to the target sequence found in the training set are classified according to their SCOP fold and Superfamily assignments. The brackets < · > denote the mean among all targets within the same cluster circled by dashed lines.

Moreover, the sequence diversity correlates positively with the contact precision at a correlation coefficient of 0.33. There are 53 cases with *T*\* < 0.4, i.e., when there is a lack of similar fold in the training set for these targets. Among them, those with high diversity, e.g., 28 targets with *N_eff_* > 1,000, have an average *L*/2 contact precision of 47% versus 30% for 25 cases with smaller sequence diversity. Likewise, for cases when similar folds are present in the training set, e.g., *T*\* > 0.6, but the *N_eff_* is low < 100, the mean contact precision is at 42%, much lower than 64%, which the mean value for cases at the same *T*\* > 0.6 criterion but with higher sequence diversity.

#### 3.2.3 Delineation of protein folds via deep-learning across SCOP folds

Notably, even when the best training structures are from a different SCOP fold, there are still many highly accurate distogram predictions as exhibited in Figure 6. Indeed, there are 102 (33%) such cases in the different fold cluster with precision > 60% among top *L*/2 predictions. Three representative examples are displayed in Figure 7. One main reason for this observation is that they may still have reasonably close or even a highly similar fold present in the training set, despite the different SCOP classifications. For 102 targets, 91/62% of them have *T*\* > 0.4/0.5, respectively. Figure 7A illustrates one such case, which is the C-terminal domain from pyruvate kinase of *Leishmania mexicana* (38). Even though there is no structure belonging to the same SCOP fold as the target, there are 12239/498/10 training structures with TM-score > 0.4/0.5/0.6 to the target structure, and the precision for contact prediction is at 98% with *d_MAE_* of 1.68 Å (*p*-value = 2.5×10^−12^) from the native distogram. A second example is delineated in Figure 7B for a domain from *Bacillus subtilis* Q45498 with unknown function. It has a *T*\* value of 0.47, among 151 training structures in the TM-score regime 0.4 to 0.5 but none higher. SAdLSA self-alignment makes a good prediction at 85% and *d_MAE_* of 2.29 Å (*p*-value = 5.1 ×10^−5)^. For these examples, it is reasonable to conceive that SAdLSA learns to predict this fold at high precision values through the training on the comparison of these structures.

**Figure 7.**
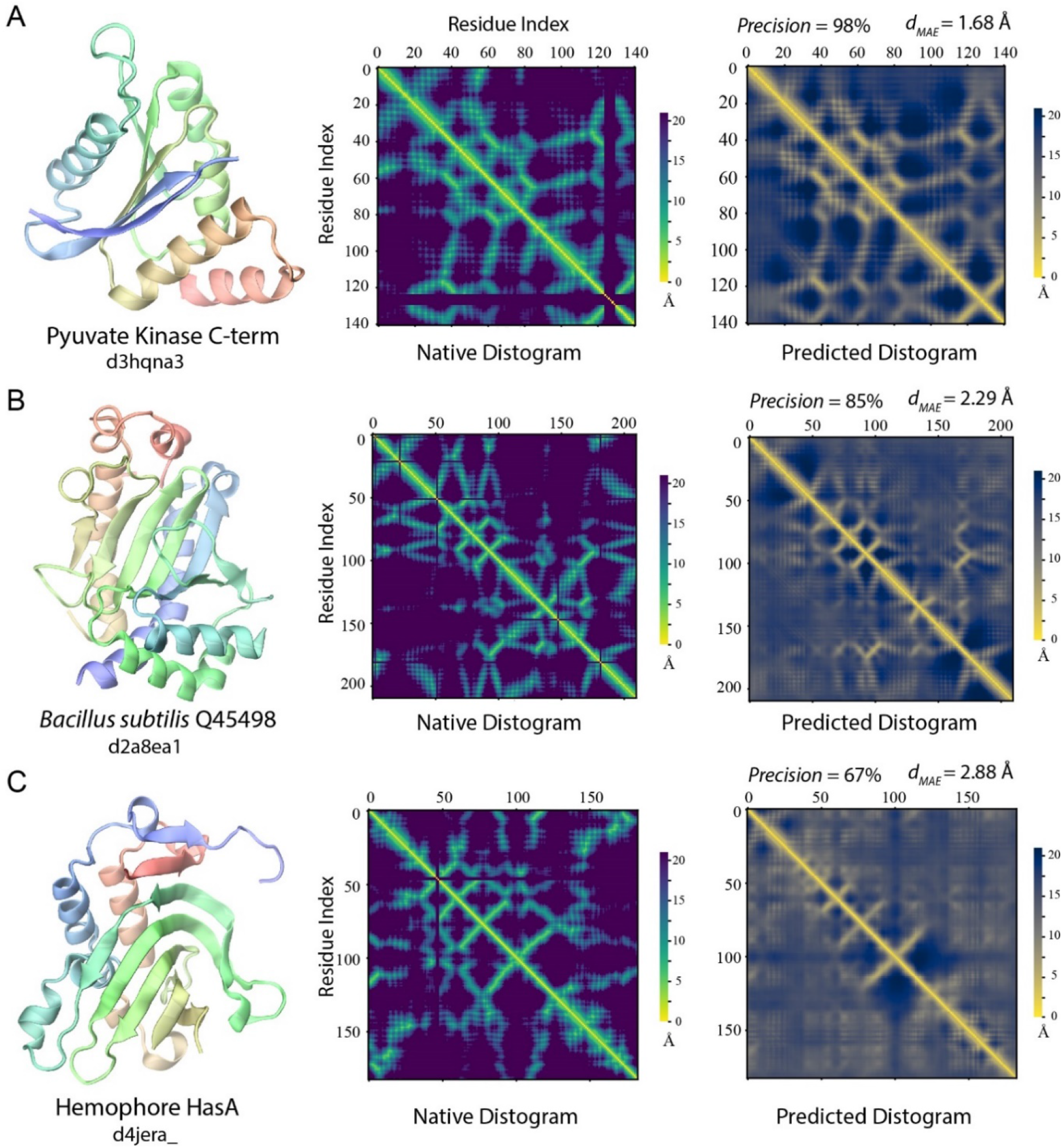
Examples of distogram predictions by SAdLSA self-alignment in comparison to native protein structures. Each panel is one example taken from targets whose fellow SCOP fold members (if any) were *not* present in the training set. The same scheme as Figure 2 was adopted to display the native structure and its distogram. Black lines in the native distograms belong to gap or non-standard amino acids in a crystal structure. The predicted distogram was calculated using the mean residue-residue distance matrix *D* formulated in the Methods. The precision values are for the medium/long-range residue contacts within top *L*/2 predictions. The value of *d_MAE_* is obtained with Eq. 2.low overall structure similarity but high similarity to some individual domains, subdomains or fragments, like the example in 3.2.1, but here it is more general and subtle without a clear definition of the domain.

In addition, there are 127 targets that do not share the same SCOP fold with any member in the training set. Yet, for 32/41 cases, SAdLSA can predict residue-residue contacts at > 50/60% precision. One such example is shown in Figure 7C. The target sequence is the hemophore HasA from *Yersinia pestis* (39). The distogram prediction via SAdLSA alignment is at 67% with a *d_MAE_* of 2.88 Å (*p*-value = 0.17). While the results are not as good as the above two cases, one can still find correct predictions between the main secondary structure elements. Here, the long-range interactions by SAdLSA are fuzzy and imprecise. Nevertheless, the result is remarkable giving that no training structure shares a TM-score > 0.4, and its *T*\* is at 0.38. In these cases, SAdLSA likely learns the packing pattern for the subdomains or fragments of a target sequence from its training structures, which may share relatively low overall structure similarity but high similarity to some individual domains, subdomains or fragments, like the example in 3.2.1, but here it is more general and subtle without a clear definition of the domain.

Lastly, one technical reason why some targets have low precision is due to the definition of this metric, which penalizes the case where very few medium or long-range contacts are present in the observed native structure, e.g., coiled-coil structures. With relatively few or even no true positives in extreme cases, the *L/n* normalization will bring down the precision value. In fact, the ratio of native contacts over the length of the sequence has the strongest correlation (0.55) with the contact precision among all five features considered. There are 22 cases whose ratio < 0.25 between the number of native medium/long-range contacts and the length of protein. Their mean precision at *L*/2 is only 9%. However, 15 of 22 have a significant *d_MAE_* < 2.68 Å (*p*-value < 0.05). Moreover, due to the lack of long-range contacts, such structures (e.g., #contacts/*L* < 0.5) are likely more flexible than compact folds, and therefore, are challenging for structure accurate prediction. In the same superfamily/different fold cluster, there are 12/49 cases forming the reddish subclusters at the edge in Figure 6. Of these 62 cases in total, 18 have a *d_MAE_* ≥ 2.68 Å. Thus, the reason for a few of these poor predictions might reflect the intrinsic propensity towards disorder for some proteins.

## 4 Discussions

What structural information does one wish to obtain from a machine-learning algorithm, given an input protein sequence? Very recently, numerous approaches have employed deep-learning techniques to predict tertiary protein structure, notably through an inter-residue distogram (2). One may argue that a more general machine-learning approach should go beyond the prediction of the tertiary structure for a single sequence to predict structural relationships between multiple protein structures, which may lead to a deeper understanding of their sequence or functional relationships. For this purpose, we introduced SAdLSA, which predicts protein sequence alignments by learning their structural alignment via deep-learning (23). In this contribution via a self-alignment analysis, we extended the previous study and explicitly demonstrate that SAdLSA learns the protein folding code. The key to understanding the deep-learning folding code lies in the analysis of the distogram prediction. Indeed, we obtain distogram predictions at surprisingly high accuracy for many folds, at a mean precision of 52% for the top *L*/2 contact predictions and a mean absolute error of 2.43 Å in inter-residue distogram predictions. In terms of *d_MAE_*, about 97% of predicted inter-residue distograms are better than expected from a background prediction, and 74% are statistically significant. This explains the advantage of SAdLSA over the classic approaches as up to a 100% improvement was observed in benchmark tests (23).

How does SAdLSA obtain its fold-depicting capability? The most important contribution comes from similar fold structures subjected to training. The algorithm was designed to pay attention to the distances between aligned residues. When these two training structures share a similar fold, the distogram of such fold can be learned, as evident in the correlation between the target and training structures (Figure 5). Additionally, if a training sequence is evolutionarily related to a target sequence, even remotely, it facilitates fold learning. More interestingly, SAdLSA can learn from training structures that go beyond the SCOP fold, e.g., cases that share no similar SCOP fold or even no similar overall fold whose TM-score > 0.4 in the training set. This seems surprising at first but may be partially explained as follows: First, a target may have many structural analogs with high structural similarity that escapes human manual classification. Second, the protein’s structure may consist of multiple-domains, subdomains, or smaller fragments whose packing pattern may be learned individually and separately from many training structures. As with all machine-learning based algorithms, since the prediction capability of SAdLSA comes from the training set, the key question is how general is the resulting model? The success in this SAdLSA self-alignment benchmark that includes many challenging cases is a good indication of its generality. But ultimately, it needs the validation in real-world, large-scale applications, ideally at the proteome-level.

Despite this success, we note that the fold-prediction ability of SAdLSA self-alignment is not as accurate as deep-learning algorithms specially designed to predict protein structures, e.g., DESTINI2. There are two main reasons for this reduced performance. First, the training of SAdLSA is on the inter-sequence, inter-residue distances between a pair of superimposed structures, rather than the intra-sequence, inter-residue distances observed in a single structure. As such, SAdLSA self-alignment performs well when a pair of structures sharing a highly similar fold are present in the training set. But this requirement is not always true, especially as our training set is derived from ~5000 SCOP domains, which is half the size of the training set for protein structure prediction, e.g., ~10000 structures for DESTINI2’s training. Only 41% of the testing targets have a close training pair at *T*\* > 0.6. Second, the current version of SAdLSA uses sequence profiles as its only input features, whereas many more features, especially the direct co-evolutionary signals that are essential for the success of DESTINI2 and the like, are not employed for a technical reason. Nevertheless, more recent deep-learning algorithms, such as AlphaFold2, directly learn from the multiple sequence alignment with an “attention” mechanism. Since the same multiple sequence alignment was used to derive the sequence profile used by SAdLSA as in DESTINI2, in principle, one may design a next generation, SAdLSA-like algorithm with a similar attention mechanism. It will not only predict protein tertiary structure via self-alignments, but also compare structures encoded by two different sequences at high precision. Such work is now underway.

## 5 Conflict of Interest

*The authors declare that the research was conducted in the absence of any commercial or financial relationships that could be construed as a potential conflict of interest*.

## 6 Author Contributions

MG and JS contributed to the conception and design of the study. MG performed the research and data analysis and wrote the first draft of the manuscript. All authors contributed to manuscript revision, read, and approved the submitted version.

## 7 Funding

This work was supported in part by the Division of General Medical Sciences of the National Institute Health (NIH R35GM118039) and the DOE Office of Science, Office of Biological and Environmental Research (DOE DE-SC0021303). The research used resources supported by the Director’s Discretion Project at the Oak Ridge Leadership Computing Facility and by the Partnership for an Advanced Computing Environment (PACE) at Georgia Tech.

## 8 Acknowledgments

We thank Bartosz Ilkowski for internal computing support and Jessica Forness for proof-reading the manuscript.

## 1 Data Availability Statement

The datasets in this study can be found at http://pwp.gatech.edu/cssb/sadlsa/.

## Notes

https://sites.gatech.edu/cssb/sadlsa/

### Competing Interest Statement

The authors have declared no competing interest.

### Summary of Updates

Minor revisions to improve the readability of the manuscript. Figures and Table are embedded in the main text.

